# The murine Microenvironment Cell Population counter method to estimate abundance of tissue-infiltrating immune and stromal cell populations in murine samples using gene expression

**DOI:** 10.1101/2020.03.10.985176

**Authors:** Florent Petitprez, Sacha Lévy, Cheng-Ming Sun, Maxime Meylan, Christophe Linhard, Etienne Becht, Nabila Elarouci, Lubka T. Roumenina, Mira Ayadi, Catherine Sautès-Fridman, Wolf H. Fridman, Aurélien de Reyniès

## Abstract

Quantifying tissue-infiltrating immune and stromal cells provides clinically relevant information for various diseases, notably cancer. While numerous methods allow to quantify immune or stromal cells in human tissue samples based on transcriptomic data, very few are available for mouse studies. Here, we introduce murine Microenvironment Cell Population counter (mMCP-counter), a method based on highly specific transcriptomic markers that allow to accurately quantify 12 immune and 4 stromal murine cell populations. We validated mMCP-counter with flow cytometry data. We also showed that mMCP-counter outperforms existing methods. We showed in mouse models of mesothelioma and kidney cancer that mMCP-counter quantification scores are predictive of response to immune checkpoint blockade Finally, we illustrated mMCP-counter’s potential to analyze immune impacts of Alzheimer’s disease. mMCP-counter is available as an R package from GitHub: https://github.com/cit-bioinfo/mMCP-counter.

## Introduction

For a large number of diseases, such as inflammatory diseases or cancer, it is often crucial to accurately determine the cellular composition of the tissue where the pathology develops, in terms of immune and stromal cell populations. An array of methods are available to obtain these data from human samples, either by immunochemistry or cytometry, or computationally from transcriptomics data^1^.

The analysis of the immune and stromal composition of tissues is particularly critical in cancer studies. Indeed, tumors are highly heterogeneous tissues which are infiltrated by a variety of immune and stromal cells^2^. It was shown that immune cell densities were associated with prognosis^3^: For instance, CD8^+^ T cells density correlates with prolonged patient survival in most cancers, whereas M2-polarized macrophages are generally associated with a poor prognosis^3^.

The transcriptome of a bulk tissue sample yields the averaged expression of genes across all the cells present in the sample. As some genes are uniquely expressed in some specific cell populations, their expression can be used to determine the abundance of the underlying cell populations. Using this property, we have previously reported on MCP-counter, a method designed to quantify the immune infiltrate of heterogeneous human tissues^4^, currently one of the best performing methods for this purpose^5^.

While murine models are widely used to decipher the pathophysiological mechanisms of various diseases, including inflammatory diseases and cancer, the computational methods currently available to measure the immune and stromal composition of murine tissues are few and limited, as compared to what is available for human samples^6^.

Here, we introduce murine Microenvironment Cell Populations counter (mMCP-counter), the adaptation of the MCP-counter method to murine samples (Figure 1), which was made possible thanks to the release of large datasets of murine sorted immune populations by the Immunological Genome Project (ImmGen)^7,8^. mMCP-counter can be accessed as an R package (https://github.com/cit-bioinfo/mMCP-counter). It takes a gene expression profiles matrix as input and returns the abundance of RNA originating from 16 defined cell populations present in the heterogeneous sample.

**Figure 1:**
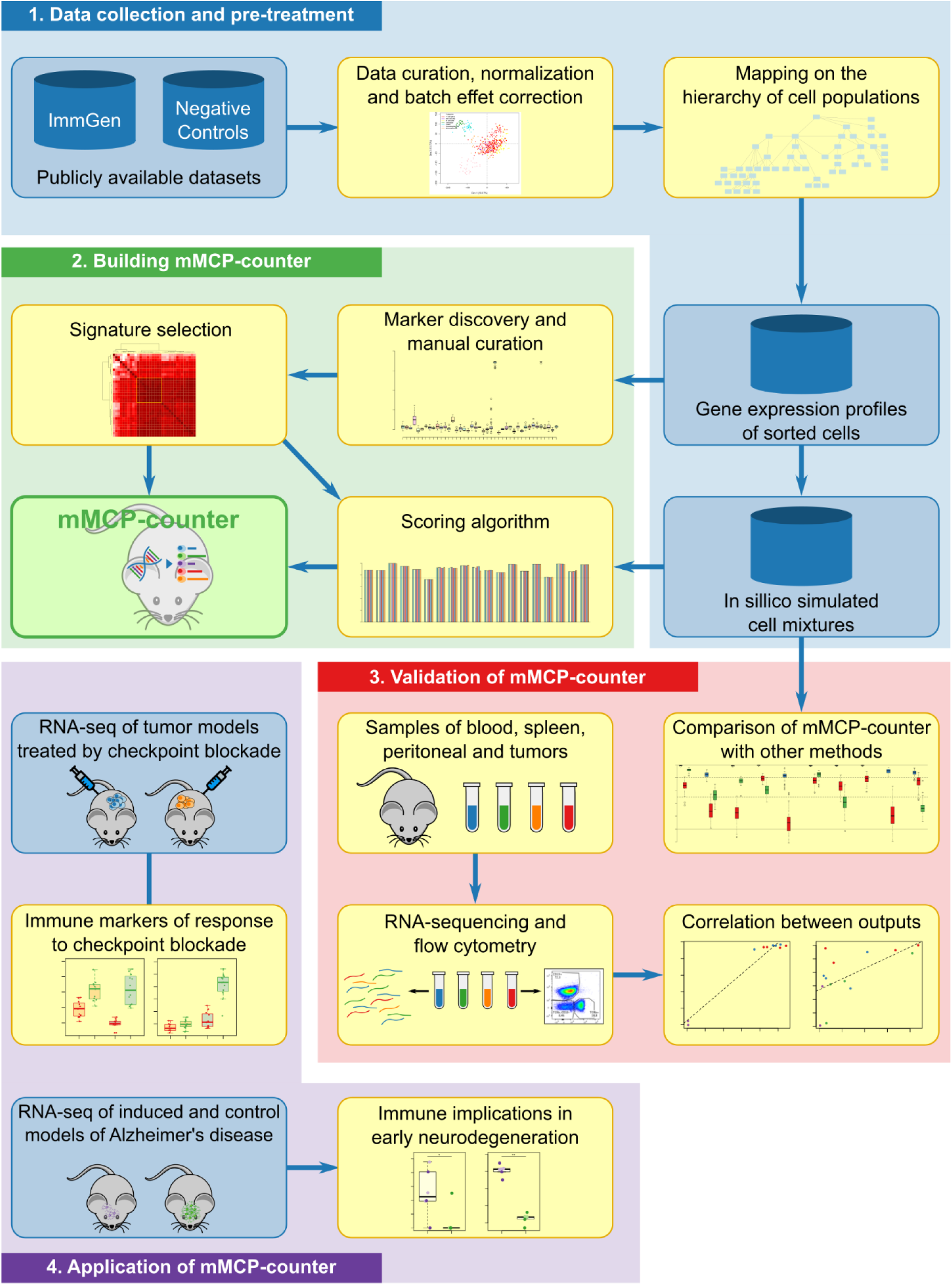
Workflow for the development, validation and application of mMCP-counter. This figure depicts 1. The data acquisition, pre-processing and normalization, as well as the mapping the cell population hierarchy; 2. The building of the methods by research and curation of cell-type specific gene signatures and optimal scoring algorithm; 3. The validation of mMCP-counter by comparison to previously published methods on simulated mixtures and by comparison to immune composition inferred by flow-cytometry; and 4. The illustration of mMCP-counter to two datasets including mouse models of kidney cancer and mesothelioma treated by immune checkpoint blockade, and murine models of early neurodegeneration in Alzheimer’s disease.

We compared the performance of mMCP-counter with other previously published methods on simulated mixtures, and we validated our approach on samples from peripheral blood, peritoneum, spleen and several grafted tumors that were analyzed by both RNA-sequencing (RNA-Seq) and flow cytometry (Figure 1). Finally, we analyzed how mMCP-counter can be used, in murine models of mesothelioma and kidney cancer, to analyze the TME differences between responders and non-responders to immune checkpoint blockade, a crucial and emerging therapy for many cancer types, and in a murine model of early neurodegeneration from Alzheimer’s disease, to identify immune and stromal cell populations in hippocampal transcriptomes.

## Results

### Prior hierarchization of cell populations

mMCP-counter relies on the identification of specific transcriptomic markers for each analyzed population. We define transcriptomic markers as having a “high” expression in a given cell population, including all its subpopulations, and “zero” expression (meaning either zero or not differentiable from the detection threshold, depending on the technologies) in any other cell population. Their detection is based on three criteria (see *signature discovery* thereafter). This approach requires to represent *a priori* both the cell categories and their inclusion relationships, as comprehensively as possible. To do so, we completed the hematopoietic tree provided by ImmGen^9^, using a survey of the literature^10,11^. The resulting hierarchy of cell populations is presented in Supplementary Figure 1a.

### Constitution of a training series

To obtain enough transcriptomic samples mapping to each of the nodes (cell categories) of the prior hierarchical model, we collected transcriptomic profiles of sorted cell populations from ImmGen. To include non-immune non-stromal negative controls parts of additional datasets were also added to our training data, including those from epithelial cells lines, hypothalamic cell lines, melanoma, pancreatic ductal adenocarcinoma and breast cancer cell lines, myoblasts and hepatocytes. Curation and normalization of data are explained in online methods.

### Signatures discovery

Only categories that were fully represented, i.e. of which all subcategories were included in the dataset, were considered for signature discovery. For each of the 55 remaining populations (Supplementary Figure 1b), all available transcripts (probes) were screened for (log2) fold-change (FC), specific fold change (sFC) and area under the ROC curve (AUC) (online methods). Features were considered as transcriptomic markers for a given population if they respected 3 criteria: FC > 2.1, sFC > 2.1 and AUC > 0.97. Figure 2a and 2b illustrate an example with probe 10442786 (*Tpsb2*), which qualified as a transcriptomic marker for mast cells. On figure 2a, we observe an overexpression of the marker in mast cells, with a log2 FC of 6.48 for the median expression as compared to the median expression in all non-mast cell samples, as well as a log2 sFC of 5.94 of the median expression in mast cells as compared to the expression variance among all non-mast cell samples. Figure 2b represents the ROC curve for this marker, showing an AUC of 0.997, thus providing sufficient sensitivity and specificity. After the automated screening, all retained transcripts were manually examined to remove transcripts that were also slightly expressed in other populations than the target populations, even though they fitted all three criteria (Supplementary Figure 2). At this step, we found transcriptomic markers for 18 denominations (Supplementary Figure 1b).

**Fig. 2:**
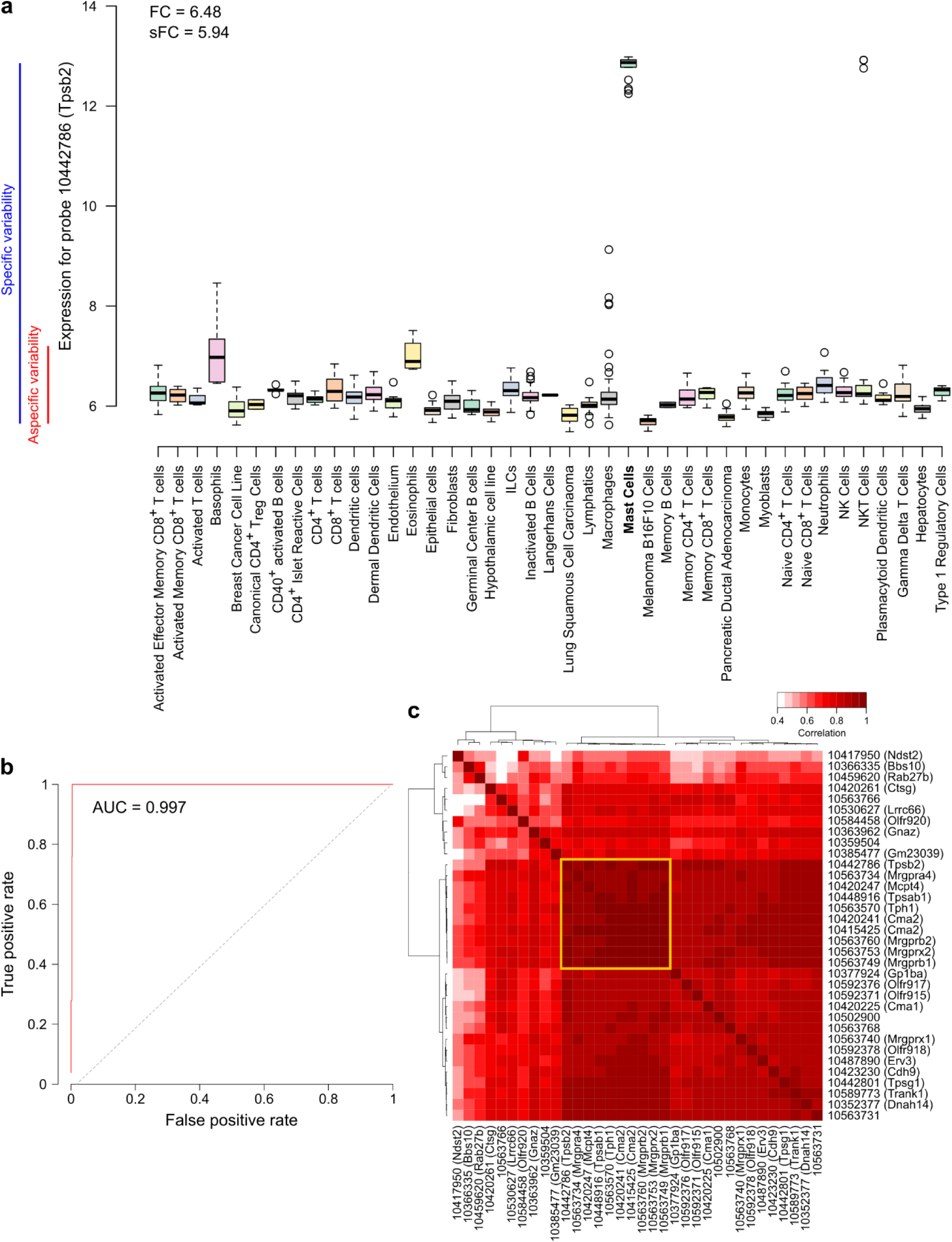
Identification of cell-type specific gene signatures: example of mast cells. **a** Expression of a transcriptomic marker (probe 10442786) of mast cells in various cell types, with the representation of the fold-change and the specific fold-change. **b** Receiver operating characteristic (ROC) curve for the same marker as in **a**. **c** Correlation heatmap of all found transcriptomic markers for mast cells. The yellow square indicates the restricted signature that was chosen for the method.

### Sub-signature selection

For some populations, a large number of transcriptomic markers was found (up to 41 for fibroblasts). A lower intra-signature correlation is likely to induce a loss of accuracy. To circumvent this potential issue, we selected a sub-signature for populations that had 8 or more markers, by choosing highest inter-correlated set of markers (Figure 2c and Supplementary Figure 3).

### Scoring metric

Given a cell population and its signature (i.e. the set of corresponding transcriptomic markers), the next step consisted in defining a metric, taking as input the expression value of this set of markers, and yielding as output a score of abundance of the cell population. To select a scoring metric, we performed tests on a dataset composed of *in silico* simulated RNA mixtures (see online methods). Six scoring metrics were considered: arithmetic mean, geometric mean, harmonic mean, quadratic mean, energetic mean and median. Each metric was tested on the mixtures by analyzing the correlations between the derived scores and the known proportions of all cell populations. The score for each scoring metric and each population are reported in Supplementary Figure 4. All metrics were found to perform similarly. The median presents the advantage of being insensitive to outliers and was therefore chosen. At this point, we discarded the dermal dendritic cells signature, as the correlation between the scores and the mixtures proportion for this population was below 0.75 (Supplementary Figure 1b).

### Ex vivo validation

To validate our approach *ex vivo*, we analyzed 14 samples of spleen (n = 4), peripheral blood (n = 4), peritoneum (n = 4) and TC1 tumors (n = 2) by flow cytometry and RNA-Seq and used the cytometry-estimated proportions of each cell type as reference (Supplementary Figure 5). We applied mMCP-counter to the RNA-seq data and computed the correlation between the flow cytometry estimates (expressed in % within living cells) and the mMCP-counter scores for hematopoietic cell populations, pooling all samples regardless of the tissue of origin (Figure 3). The signature for canonical CD4^+^ regulatory T cells failed this validation step (Figure 3b and Supplementary Figure 1b). However, for all other available populations (Figure 3a), there was a good agreement between the mMCP-counter scores and the proportions obtained by flow cytometry, with correlation comprised between 0.629 (eosinophils) and 0.975 (CD8^+^ T cells).

**Figure 3:**
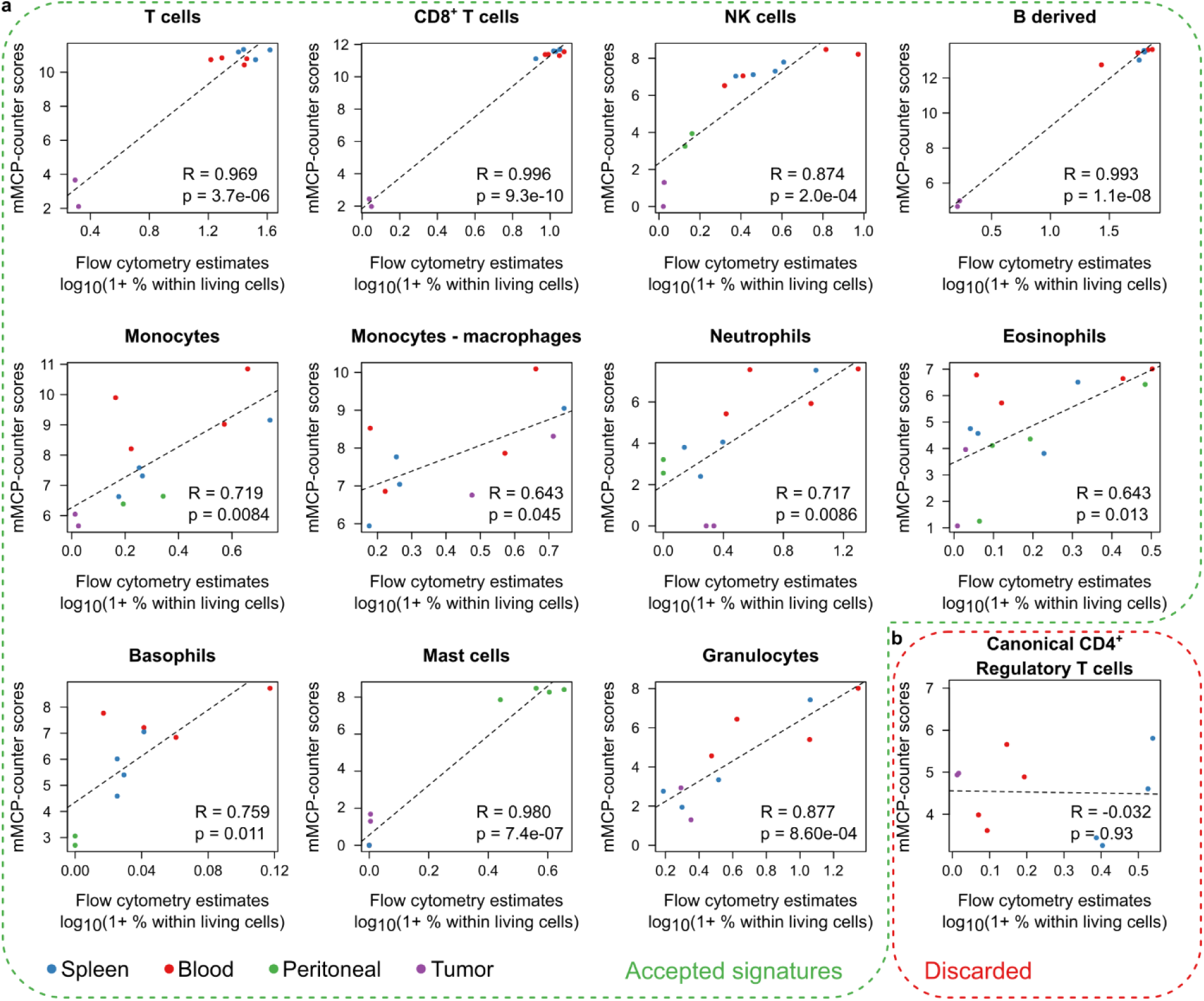
Validation of mMCP-counter on *ex vivo* data by comparison to flow cytometry data. **a** Correlation graphs between the flow cytometry estimates (logarithmic scale, expressed in percent of the total of living cells) and the mMCP-counter scores for populations for which the signature was accepted. Each graph corresponds to a different population. The dotted line shows the linear regression model. Correlations are estimated with the Pearson correlation. **b** Correlation graph for canonical CD4^+^ regulatory T cells, presented as in **a**. Following this validation step, this signature was discarded.

### Comparison with other published methods

Other methods have been previously reported to analyze the composition of heterogeneous samples in murine models^12,13^. DCQ (Digital Cell Quantification) is an algorithm that, given gene expression data and prior knowledge on immune cell types transcriptomic profiles returns the cell abundances for a wide variety of immune cells^12^; it was designed using RNA-Seq data. ImmuCC^13^ is derived from the method proposed by CIBERSORT^14^ and adapted by finding markers for murine populations; it was first designed for micro-arrays, but an updated version is adapted for RNA-Seq data^15^. We applied both methods on 50 sets of *in silico* RNA mixtures (each with 50 samples, see online methods) to assess each method’s performance on cell subtypes that were quantified by both mMCP-counter and another method (Figure 4). On all considered populations, mMCP-counter outperformed both ImmuCC and DCQ (p < 3e-13 for all comparisons). mMCP-counter also proved to consistently perform well, with the median correlation between mixture compositions and scores above 0.8 for all considered populations, whereas as ImmuCC and DCQ performance greatly varied depending on the populations.

**Fig. 4:**
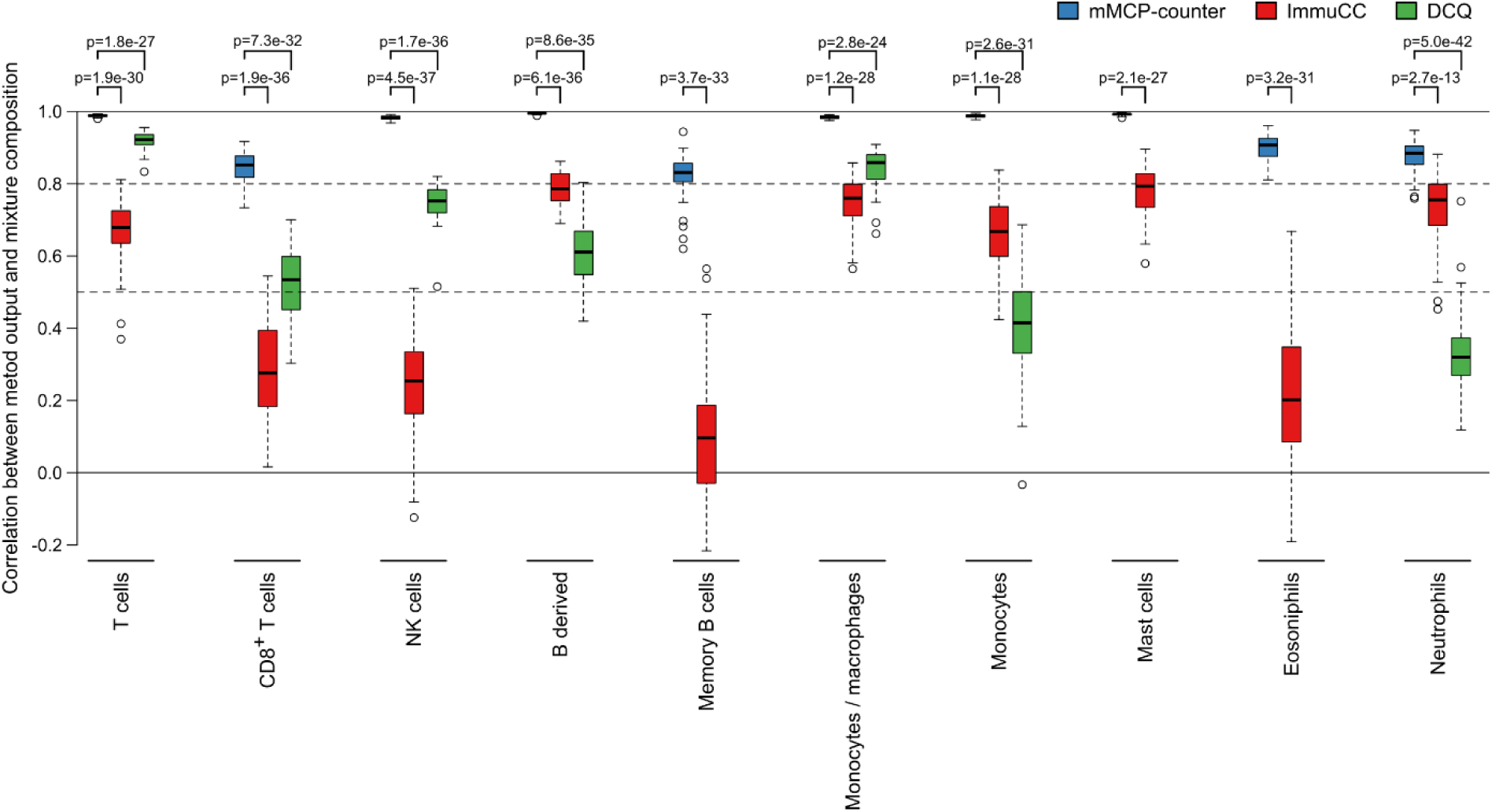
Comparison of the performance of mMCP-counter with other published methods. The three methods have been applied to 50 simulated RNA mixture sets, each comprising 50 randomized mixtures and this graph shows the Pearson correlation between the mixture compositions and the scores returned by the methods for each population on all mixture sets. Full lines indicate correlation equal to 0 and 1. Dashed lines indicate correlations equal to 0.8 and 0.5.

### mMCP-counter discriminates tumor types and responders to immune checkpoint blockade

Immune checkpoint blockade (ICB) has become in the last decade a crucial treatment option for cancer patients. The response rate to such drugs strongly varies depending on the malignancy, and identifying patients likely to respond remains a challenge. Mouse pre-clinical models greatly help to identify markers of response that are potentially useful in human clinical trials. We therefore applied mMCP-counter to pre-treatment samples of mouse models of kidney cancer and mesothelioma that have been treated with a combination of CTLA-4 and PD-L1 blockade^16^. We could therefore investigate whether mMCP counter could detect differences in the tumor micro-environment (TME) composition between cancer types and between tumors responding or not to ICB. An unsupervised analysis (Figure 5a) revealed that the TME, as analyzed by mMCP-counter, principally discriminates malignancies based on the tumor type. Within each tumor type, the unsupervised clustering on the mMCP-counter scores allows to discriminate between two groups associated with response to ICB, suggesting that the TME composition is tightly associated with response to ICB.

**Figure 5:**
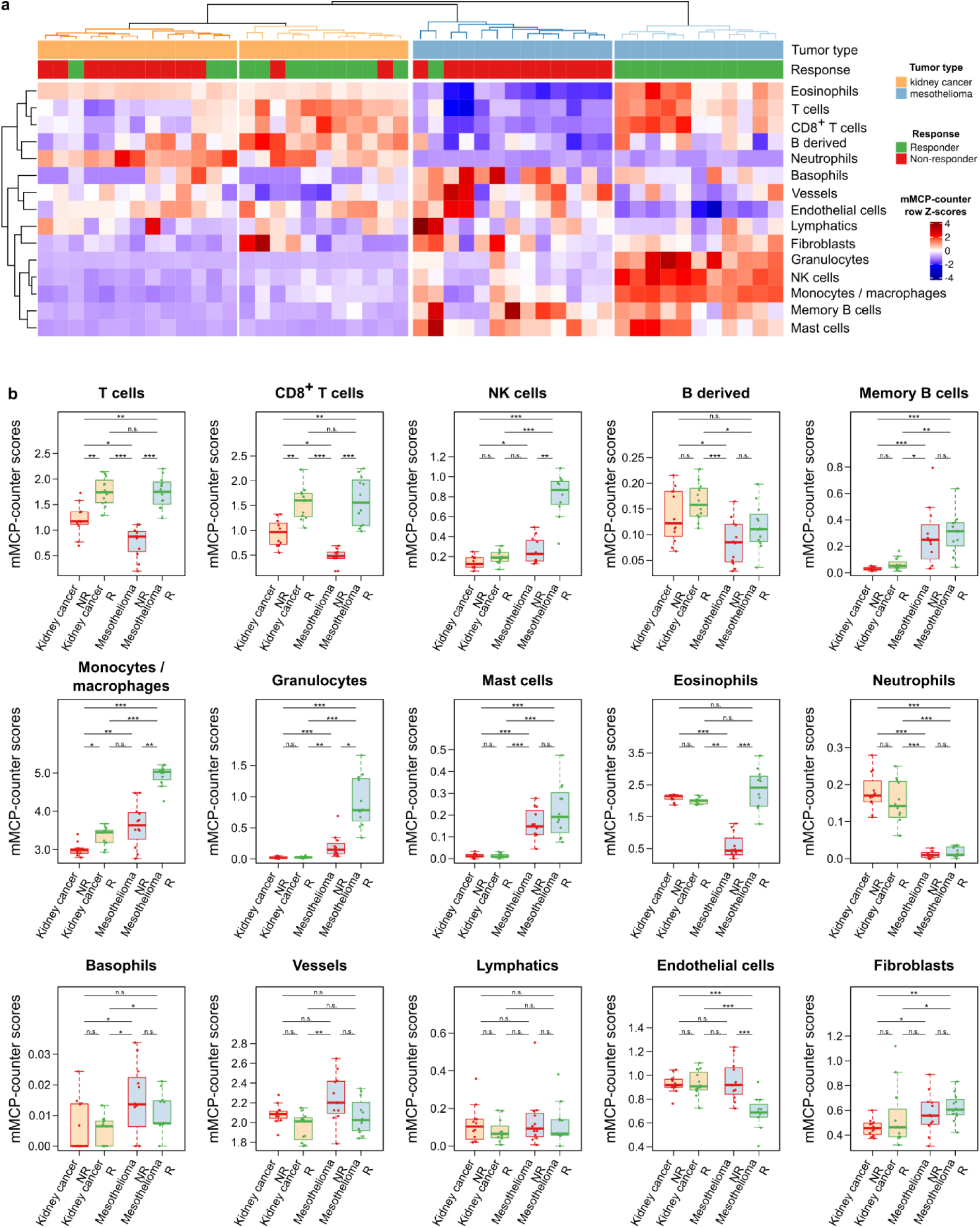
mMCP-counter allows to discriminate between tumor types and between responders and non-responders to immune checkpoint blockade. **a** Heatmap showing that clustering of tumors on mMCP-counter scores accurately separates tumors based on the tumor type (first line) and the response to immune checkpoint blockade (second line). The heatmap illustrates row Z-scores for all included cell populations. **b** Detailed differences in mMCP-counter scores between responders and non-responders to ICB in both kidney cancer and mesothelioma models. Comparisons are computed using Kruskal-Wallis tests followed by post-hoc Dunn test for pairwise comparisons, with Benjamini-Hochberg correction for multiple testing. * p<0.05, ** p<0.01, *** p<0.001, n.s. p≥0.05.

In more details, we also analyzed the association between the scores for each population and response, in both models (Figure 5b). This revealed associations with response that are found in both kidney cancer and mesothelioma models. Indeed, the two models showed that responsive tumors had an increased infiltration by T cells, CD8+ T cells, monocytes / macrophages as compared to tumors that resisted the ICB treatment. However, other TME differences between responders and non-responders appear to be cancer type-specific. Indeed, in mesothelioma, responders exhibited more NK cells, granulocytes, eosinophils, and less endothelial cells than non-responders, while these associations were not found in kidney cancer models. Moreover, some populations were particularly differentially present in the two tumor types, including NK cells, B derived cells, memory B cells, monocytes/macrophages, granulocytes, mast cells, neutrophils, basophils and fibroblasts.

### mMCP-counter identifies immune and stromal correlates of early Alzheimer’s disease onset

Alzheimer’s disease (AD) has been modelled by a bitransgenic mouse model called CK-p25 which can overexpress p25 when induced through the calcium/calmodulin-dependent protein kinase II (CK) promoter, as compared to CK control mice^17^. p25 triggers an aberrant activation of cyclin-dependent kinase 5, which in turn increases phosphorylation of pathological substrates including tau. We obtained RNA-seq data from the hippocampus of CK-p25 mice, thereafter labeled as AD mice, at 2 or 6 weeks into neurodegeneration, as well as similar data from control CK mice^18^. Using mMCP-counter, we observed that the immune and stromal composition of the samples neatly segregated mice with AD from CK mice (Figure 6a), suggesting that AD impacts the hippocampus’ immune infiltration and vascularization.

**Figure 6:**
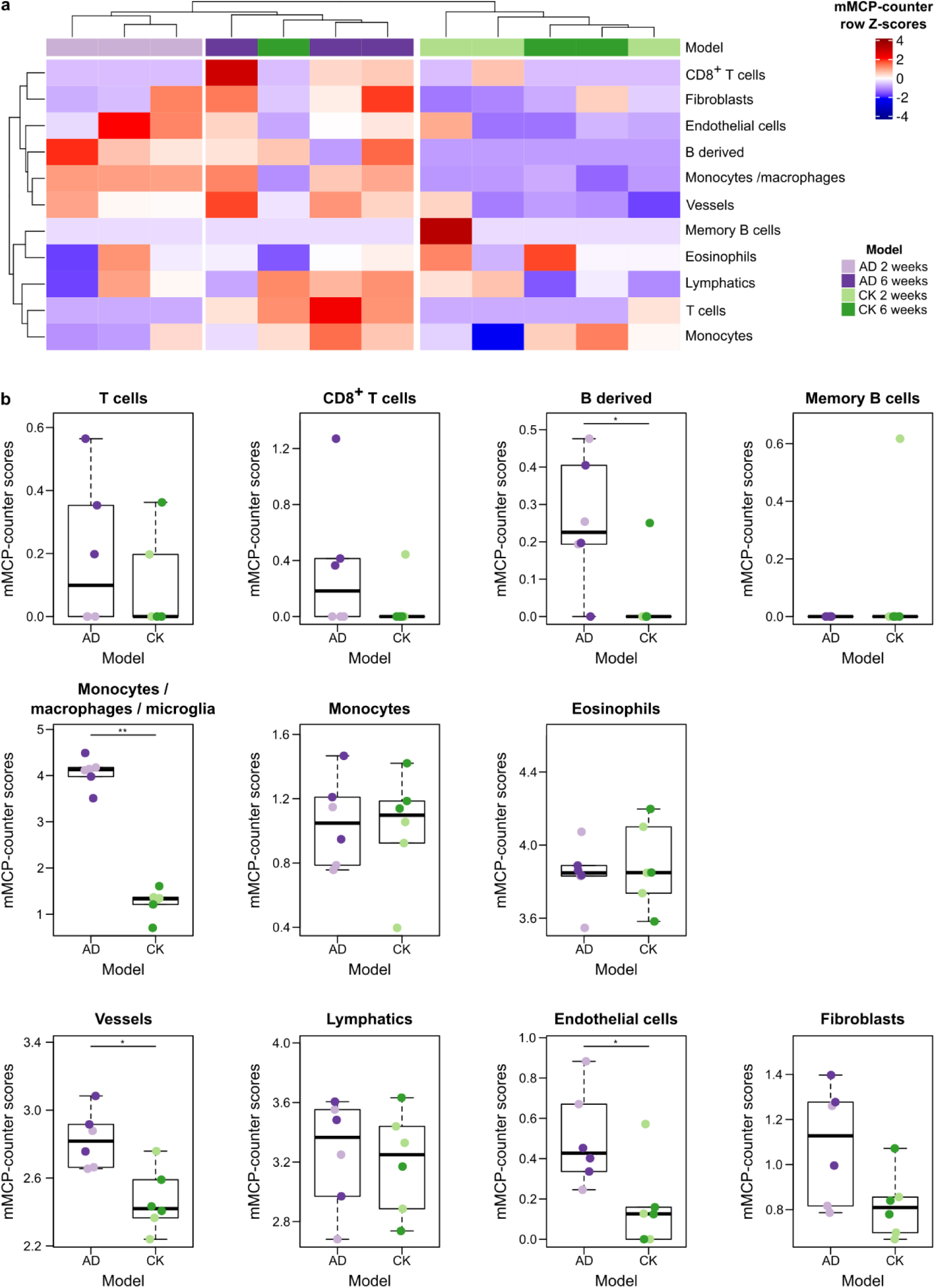
mMCP-counter allows to discriminate between control CK mice and Alzheimer’s disease brain tissues. **a** Heatmap showing that clustering of samples on mMCP-counter scores accurately separates hippocampus samples from control CK samples and induced Alzheimer’s disease (AD) at different time points. The heatmap illustrates row Z-scores for all included cell populations. **b** Detailed differences in mMCP-counter scores between CK and induced AD hippocampus samples. The colour code of the individual data points refers to the legend of panel **a**. Comparisons are computed using Mann-Whitney tests. * p<0.05, ** p<0.01.

In details, we notably observed that AD mice hippocampus had a higher infiltration by B derived cells, but not memory B cells, more macrophages (since they have an increased score for monocytes / macrophages, but not monocytes), and more endothelial vessels (Figure 6b). The increase in macrophages is likely due to microglia, the resident macrophages of the central nervous system. Indeed, using single-cell RNA-sequencing data from the *Tabula Muris* project^19^, we noticed a strong expression of the Monocytes / macrophages signature by microglial cells in the brain (Supplementary Figure 6). No significant alterations were found for T cells, eosinophils, lymphatics and fibroblasts. We noted that 6-week AD mice showed an increased presence of CD8^+^ T cells in the hippocampus, although the limited number of mice did not allow to reach significance (p=0.01).

## Discussion

Here we introduce mMCP-counter, a method to quantify immune and stromal cell populations in heterogeneous murine samples. mMCP-counter is based on the identification of highly specific transcriptomic signatures for each of the cell populations considered. We found robust signatures for a total of 16 populations: 12 immune populations (T cells, CD8^+^ T cells, NK cells, B derived cells, memory B cells, monocytes/macrophages, monocytes, granulocytes, mast cells, eosinophils, neutrophils and basophils) as well as 4 stromal populations (vessels, lymphatics, endothelial cells and fibroblasts).

We validated mMCP-counter method by comparing the scores with flow cytometry estimates on blood, spleen, peritoneal and tumor samples and demonstrate a strong correlation between both methodologies. We also showed that mMCP-counter allows a significant improvement over the existing methods by comparing its performance to previously developed methods on large simulated datasets.

Applied to mouse models of kidney cancer and mesothelioma treated by combination of immune checkpoint blockade therapies, mMCP-counter allows to decipher the differences of tumor microenvironment composition between both tumor types and between responders and non-responders to immune checkpoint blockade. Alongside known associations between TME composition and response to ICB, such as T cells and CD8^+^ T cells, mMCP-counter revealed that tumors responsive to ICB had a higher infiltration by monocytes and/or macrophages. Moreover, malignancy-specific differences between responders and non-responders could be observed, that were not previously reported. Finally, there were strong differences in the overall composition of the TME between mesothelioma and kidney cancer models. Due to the rapidly increasing, almost impossible to handle, number of agents tested in immunotherapy clinical trials^20,21^, it is of paramount importance to test them in pre-clinical models with a method that robustly and sensitively quantifies the TME composition. Altogether, mMCP-counter may help finding the rationale for potential cancer-specific combination strategies and drive more efficient personalized cancer medicine.

To assess the applicability of mMCP-counter beyond the field of cancer, we analyzed a model of Alzheimer’s disease. Thus, we applied mMCP-counter to hippocampal transcriptomics data from a murine model of Alzheimer’s disease, comparing mice with induced AD with controls. The most striking difference was an increased expression of the monocyte / macrophage signature in AD mice. This may be explained by the detection by mMCP-counter of microglia, which have been shown to be a prominent marker of AD^22^. Conversely, the role of B cells is disputed and looked as inessential in AD^22^. Here, we also noticed an increased in B lineage cells. Finally, we also noted an increased presence of endothelial vessels in AD mice, in line with reports of angiogenesis in AD^23^. mMCP-counter could help further analyze the immune and stromal impacts of AD and other neurodegenerative syndromes. Although it did not reach significance, a trend towards an increase in the presence of CD8^+^ T cells in the hippocampus of AD mice between weeks 2 and 6 was observed. This concords with recent observation of presence of CD8+ T cells in the hippocampi of AD human patients^24^.

mMCP-counter is fast and memory-efficient to compute the scores. The abundance scores it provides are shown to be linearly related to the known abundances of the related cell populations in validation data. These scores can thus safely be compared across the samples of a given series, as illustrated here in two examples. mMCP-counter scores are given in population-dependent arbitrary units. As such, the intra-sample ratio of the scores of two distinct cell populations is not an accurate estimate of the actual intra-sample ratio of these two populations, as reported for MCP-counter^25^. However, such a ratio could still be compared across samples within a series, as by construction it would also be linearly correlated to true ratios. This is a major difference with some methods, including CIBERSORT and its derivatives such as ImmuCC, which instead enable intra-sample comparison but do not allow inter-sample comparisons^5,25^.

For human samples, a large number of methods are available and are part of a global set of methods to study cancer immunity^26^. However, they differ in their performance, and signatures appear to be the most critical aspect of such approaches^27^. The robust and stringent definition of signatures of MCP-counter allows it to be among the best performing ones^5^. However, only few of these methods were available for murine models to this day. Here, we have kept the same methodology to define gene signatures that are highly specific for the considered cell populations, which could explain that mMCP-counter outperforms the other approaches. Indeed, to build mMCP-counter, we chose to use very stringent definitions of specific transcriptomic markers. This allows a precise estimation of all measured populations, but it is at the expense of the number of populations that can be accounted for. Indeed, other methods, such as DCQ estimate changes in more than 70 immune populations, where we only quantify 12, plus 4 stromal populations. mMCP-counter therefore cannot estimate all precise functional orientations but outperforms DCQ for the populations where it applies. Nevertheless, robust signatures for memory B cells were identified here. Similarly, there is a growing interest in the heterogeneity of cancer-associated fibroblasts^28,29^. To include more details as to fibroblasts subtypes in mMCP-counter would have required far more detailed data on sorted fibroblasts subtypes than what is currently available.

mMCP-counter can provide extremely useful information in cancer murine models. A major asset of mMCP-counter, compared to previously reported methods, is that it allows to simultaneously study immune and stromal cell populations. The clinical relevance of the microenvironment composition is not only documented for immune cells^3^, but also for stromal cells: blood vessels and angiogenesis are key players in cancer development and metastasis^30^; lymphatic vessels are associated with metastasis^31^ and the impact of fibroblasts raises a growing interest^32^.

Beyond the field of cancer, mMCP-counter may also have broad applications in murine models of diseases in which immunity, inflammation or angiogenesis play crucial roles. In particular, it can be applied to models of neurodegenerative syndromes such as Alzheimer’s disease, well-known auto-immune^33^ and inflammatory^34,35^ diseases, but also atherosclerosis^36^.

## Online Methods

### Data accession, curation and normalization

Data included in the discovery dataset included the two micro-array releases of ImmGen (Gene Expression Omnibus (GEO) accession numbers GSE15907 and GSE37448), and parts of several datasets for: epithelial cells (GSE27456 and GSE74317), breast cancer (GSE25525, GSE54626 and GSE78698), hypothalamic cell line (GSE61402), melanoma B16F10 cells (GSE84155), pancreatic ductal adenocarcinoma (GSE48643), myoblasts (GSE26764) and hepatocytes (GSE18614). Raw CEL files were used and data was normalized through frozen robust multi-array analysis^37^ with the R package fRMA. Batch effect was correcting using ComBat^38^ from the R package sva. Consistency within the data was verified using principal components analysis with the R package FactoMineR^39^ and outliers were discarded.

Application data were downloaded from GEO (accession numbers GSE93017 and GSE117358). For GSE117358, data was normalized at the 75^th^ percentile of gene expression.

### Signatures discovery

For signatures discovery, only populations for which all subsets were present in the dataset and that have appropriate negative samples were taken into account. all probes were screened for (log2) fold-change (FC), specific fold change (sFC) and area under the ROC curve (AUC). FC and sFC are defined as follows:

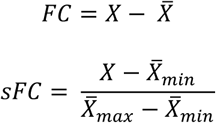

where *X* is the centroid (i.e. the mean over all samples) for the population of interest, 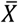 the centroid of all other samples, and 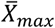 and 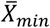, denote, respectively, the maximum and the minimum of cell-type-specific centroids for population different from the population of interest.

Signatures were built using the following cut-offs: FC > 2.1, sFC > 2.1 and AUC > 0.97. After this automated screening, all retained probed were manually curated to verify the accuracy of the selection. All signatures that contained more than 8 putative transcriptomic markers underwent an additional selection process. A sub-signature with strong inter-marker correlation was kept following hierarchical clustering of the whole signature transcriptomic markers. The hierarchical clustering was made using R, with Euclidian metric and Ward’s linkage criterion.

### In sillico mixtures preparation

The *in sillico* simulated mixtures were computed as follows: firstly, weights for all included populations were chosen randomly. Pure transcriptomic profiles for all populations were computed with the expression of all genes being the mean expression over all the corresponding samples in the ImmGen dataset. Finally, the mixture transcriptome was computed as follows:

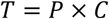

where *T* is the transcriptomic matrix with genes in lines and samples (mixtures) in columns, *P* is the pure profiles matrix with genes in lines and cell populations in columns and *C* is the mixture composition matrix, with populations in lines and samples (mixtures) in column, the sum of each column being equal to 1.

To evaluate the various scoring algorithms, 24 mixtures were simulated with random proportions of each cell populations. For the comparisons between mMCP-counter and other methods, 50 sets of mixtures were selected. For each mixture set, the random proportions for 50 mixtures were simulated using a Dirichlet distribution with shape parameters 1 (Activated Effector Memory CD8 T cells), 1 (Activated Memory CD8 T cells), 1 (Activated T cells), 2.5 (CD4 T cells), 2.5 (CD8 T cells), 1 (Memory CD4 T cells), 1 (Memory CD8 T cells), 1 (Naïve CD4 T cells), 1 (Naive CD8 T cells), 0.5 (Gamma delta T cells), 5 (NK cells), 5 (CD40 activated B cells), 2.5 (Germinal center B cells), 2.5 (Inactivated B cells), 2.5 (Memory B cells), 5 (Monocytes), 5 (Macrophages), 2.5 (Mast cells), 2.5 (Eosinophils), 2.5 (Neutrophils) and 2.5 (Basophils).

### Comparison with other published methods

mMCP-counter, DCQ and ImmuCC were run independently on each of the 50 sets of 50 mixtures defined above, aggregated by gene. The ImmuCC algorithm and signature matrix was accessed on GitHub (https://github.com/chenziyi/ImmuCC) and ran on the mixtures locally. DCQ was run using the dcq function from the ComICS R package. All methods were run using default parameters. To account for populations similar to mMCP-counter, scores of all subpopulations were summed.

### Animal experiment

Eight to ten-week-old female C57/BL6 mice were purchased from Charles River Laboratories. The use of animals was followed the institutional guidelines and the recommendations for the care and use of laboratory animals with approvals APAFIS#34\0-2016052518485390v2 and #9853-2017050211531651v5 by the French Ministry of Agriculture. Mice were sacrificed and spleens were surgically removed and were pressed through a 70 µm cell strainer (BD Falcon) for single cell suspension preparation. Blood was obtained with cardiac puncture or from submandibular vein. Peritoneal cells were obtained by washing peritoneal cavity with 3 – 4 ml of PBS twice. Red blood cells were lysed by ACK lysing buffer and cells were then washed with PBS with 2% of fetal bovine serum (FBS). All cells were resuspended in ice cold PBS with 2% FBS for FACs staining.

TC-1, tumor cells derived from mouse lung epithelial cells and transformed by human papillomavirus^40^ were cultured *in vitro*. Cells were washed with PBS and 4 × 10^5^ cells were inoculated subcutaneously in the right flank with 200 µl PBS. Twenty-six days later, tumors were surgically removed and single cell suspension is prepared for FACs analysis.

### Flow cytometry

For flow cytometry, cells were stained with the following monoclonal antibodies: PE-conjugated anti-CD4, Fitc-conjugated anti-CD8, BV786-conjugated anti-CD11c, PE-Cy7 conjugated anti-CD45, BV605 conjugated anti-NK-1.1 (all from BD Biosciences), eFluor 450-conjugated anti-CD11b, Alexa Fluor 700-conjugated anti-CD19, APC-eFluor 780-conjugated anti-CD19, PerCP-eFluor 710-conjugated anti-CD49b, PE-CF594-conjugated anti-Siglec-F, (all from eBioscience), Brilliant Violet 785-conjugated anti-CD11b, APC/Fire 750 conjugated anti-TCR-β, Pacific Blue conjugated anti-GL7, Fitc-conjugated anti-FcεRIa, Alexa Fluor 700-conjugated anti-F4/80, Brilliant Violet 605-conjugated anti-Ly-6C and Brilliant Violet 650-conjugated anti-Ly-6G (all from BioLegend). Cells were stained for 30 min in the dark at 4°C and were washed with PBS with 2% of FBS. For Foxp3 staining, cells were fixed and permeabilized with eBioscience Foxp3/Transcription Factor Staining Buffer Set according to manufacturer’s protocol (eBioscience). All stainings were done with Fc block (BD Biosciences). Cells were then analyzed on a BD LSRFortessa cell analyzer (BD Bioscience). Flow cytometry data analysis was performed using Flowjo analysis platform (FlowJo, LLC).

Living cells were identified by LIVE/DEAD Fixable aqua dead cell stain kit (ThermoFisher Scientific) and singlet cells were gated before further analysis. T cells are identified as CD19^-^B220^-^CD11b^-^NK1.1^-^TCRβ^+^ cells, CD8^+^ T cells are CD19-B220-CD11b-NK1.1-TCRβ^+^CD8^+^ cells, T_reg_ cells are CD19-B220-TCRβ^+^CD4^+^CD25^+^Foxp3^+^ cells, NK cells are CD19^-^TCRβ^-^NK1.1^+^ cells, B derived cells are CD19^+^B220^+^TCRβ^-^ cells, memory B cells are CD19^+^B220^+^CD38^+^CD80^+^IgD^lo^ cells, neutrophils are CD19^-^TCRβ^-^CD11c^-^CD11b^+^Ly6G^+^ cells, eosinophils are CD19^-^TCRβ^-^CD11c^-^CD11b^lo^Ly6G^-^SiglecF^+^ cells, basophils are CD19^-^TCRβ^-^CD11c^-^CD11b^+^Ly6G^-^CD117^-^FcεRIa^+^CD49d^+^ cells, mast cells are CD19^-^TCRβ^-^CD11b^-^FcεRIa^+^CD117^+^ peritoneal cells, monocytes are CD19^-^TCRβ^-^CD11c^-^F4/80^-^ CD11b^+^CD115^+^ cells, macrophages are CD19^-^TCRβ^-^F4/80^+^CD11b^+^ peritoneal cells.

### RNA preparation

Splenocytes, peripheral blood cells, peritoneal cells and tumor cells were washed with PBS and were counted. Cells were centrifuged at 400 *g* for 5 min and supernatant was removed. Cells pellet (< 3 × 10^6^ cells) was resuspend in 350 µl of RLT buffer (Qiagen). RNA was extracted with RNeasy Mini kit (Qiagen) according to manufacturer’s protocol.

### RNA sequencing

mRNA library preparation was realized following manufacturer’s recommendations (KAPA mRNA HyperPrep ROCHE). Library purity/integrity were assessed using an Agilent 2200 Tapestation (Agilent Technologies, Waldbrunn, Germany). Final 7 samples pooled library prep were sequenced on Nextseq 500 ILLUMINA with MidOutPut cartridge (2×130Millions of 75 bases reads), corresponding to 2×18Millions of reads per sample after demultiplexing.

### Analysis of single-cell RNA-seq data

The analysis of the single-cell RNA-seq data from the *Tabula Muris* consortium^19^ was accessed on the Single Cell Expression Atlas^41^ from EMBL-EBI at https://www.ebi.ac.uk/gxa/sc/home on March 4th, 2020 and analyzed online on this platform.

### Statistical analysis

All statistical analyses were made using R 3.6.2 with packages gtools, ComICS and ComplexHeatmap^42^. Pearson’s correlation was used to compare two quantitative variables. The comparisons between mMCP-counter and other methods (figure 4) were assessed using t tests. For other comparisons between a quantitative variable and a 2-level qualitative variable, we used Mann-Whitney tests. For 3 or more levels, we used Kruskal-Wallis with post-hoc Dunn test for pairwise comparisons with Benjamini-Hochberg correction. Associations between two quantitative variables were assessed using Pearson correlation and its correlation test.

## Acknowledgements

The authors thank Robert D. Schreiber for insightful discussions. This work was supported by the Institut National de la Santé et de la Recherche Médicale, the Université de Paris, Sorbonne University, the Programme Cartes d’Identité des Tumeurs (CIT) from the Ligue Nationale Contre le Cancer, grants from Institut National du Cancer (HTE-INSERM plan cancer, C16082DS), Cancer Research for Personalized Medecine programme (CARPEM T8), “FONCER contre le cancer” programme and Labex Immuno-Oncology (LAXE62_9UMRS972 FRIDMAN). This work benefited from equipment and services from the iGenSeq core facility, at Institut du Cerveau et de la Moelle épinière. This project has received funding from the European Union’s Horizon 2020 research and innovation programme under grant agreement No 754923. The materials presented and views expressed here are the responsibility of the authors(s) only. The EU Commission takes no responsibility for any use made of the information set out. FP was supported by CARPEM doctorate fellowship.

## Author contributions

FP, WHF, AdR and CSF designed the study. CL and NE accessed and curated publicly available datasets. FP, SL, MM, CL and EB analyzed the data. CMS and LTR carried the murine and flow cytometry experiments and RNA extraction. MA designed the RNA-seq protocol. FP, WHF, AdR, CSF and MM wrote the manuscript, which was reviewed and approved by all authors.

