## Supplementary figures for "The murine Microenvironment Cell Population counter method to estimate abundance of tissue-infiltrating immune and stromal cell populations in murine samples using gene expression"

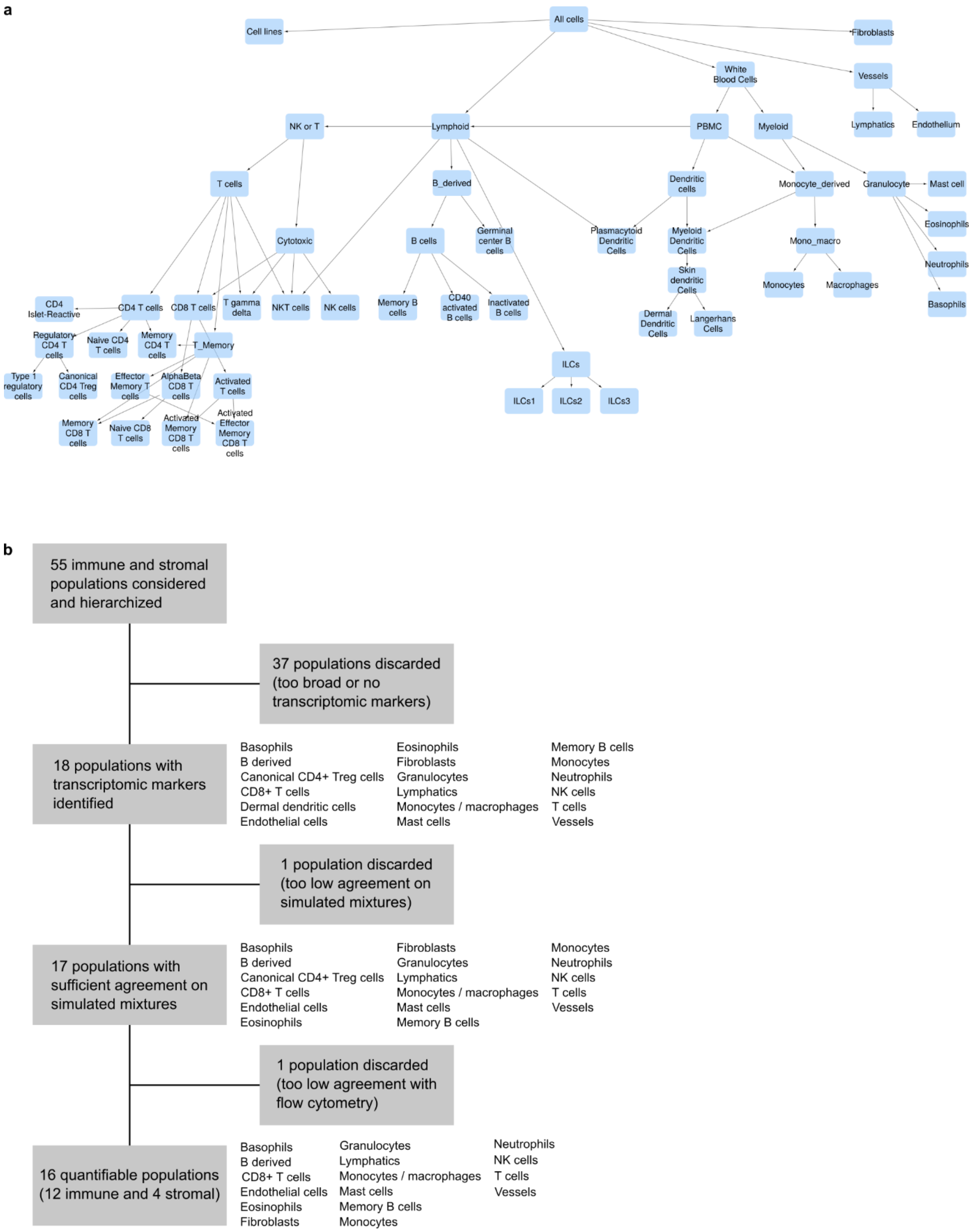


**Supplementary Figure 1: Cell populations and signatures**

**a** Cell populations hierarchy used in the present study. **b** Counts of the number of signatures identified and discarded during the various validation steps.

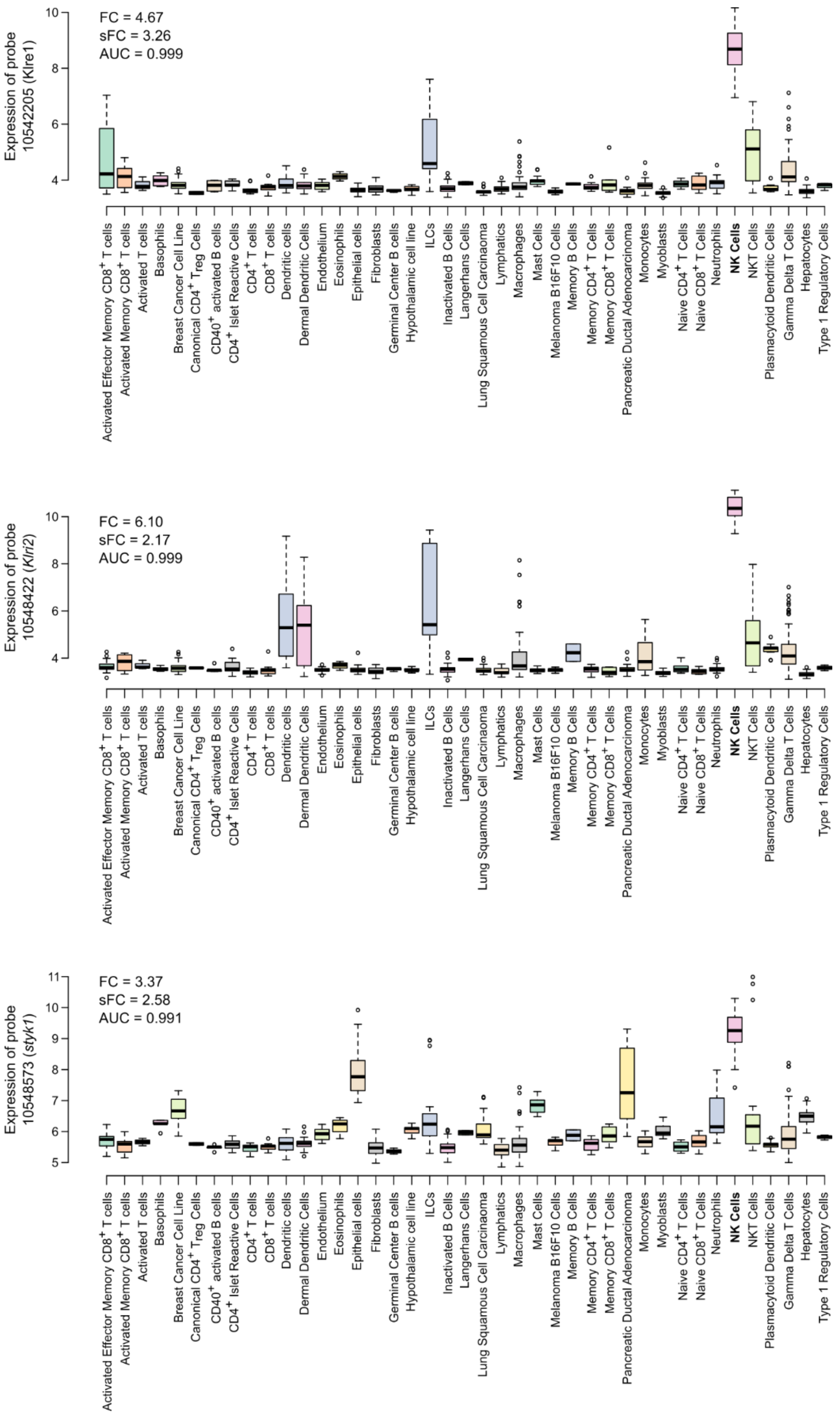


**Supplementary Figure 2: Examples of discarded transcriptomic markers**

The three markers presented here satisfied the criteria to be considered as transcriptomic markers for NK cells, but they were discarded during manual curation due to insufficient specificity.

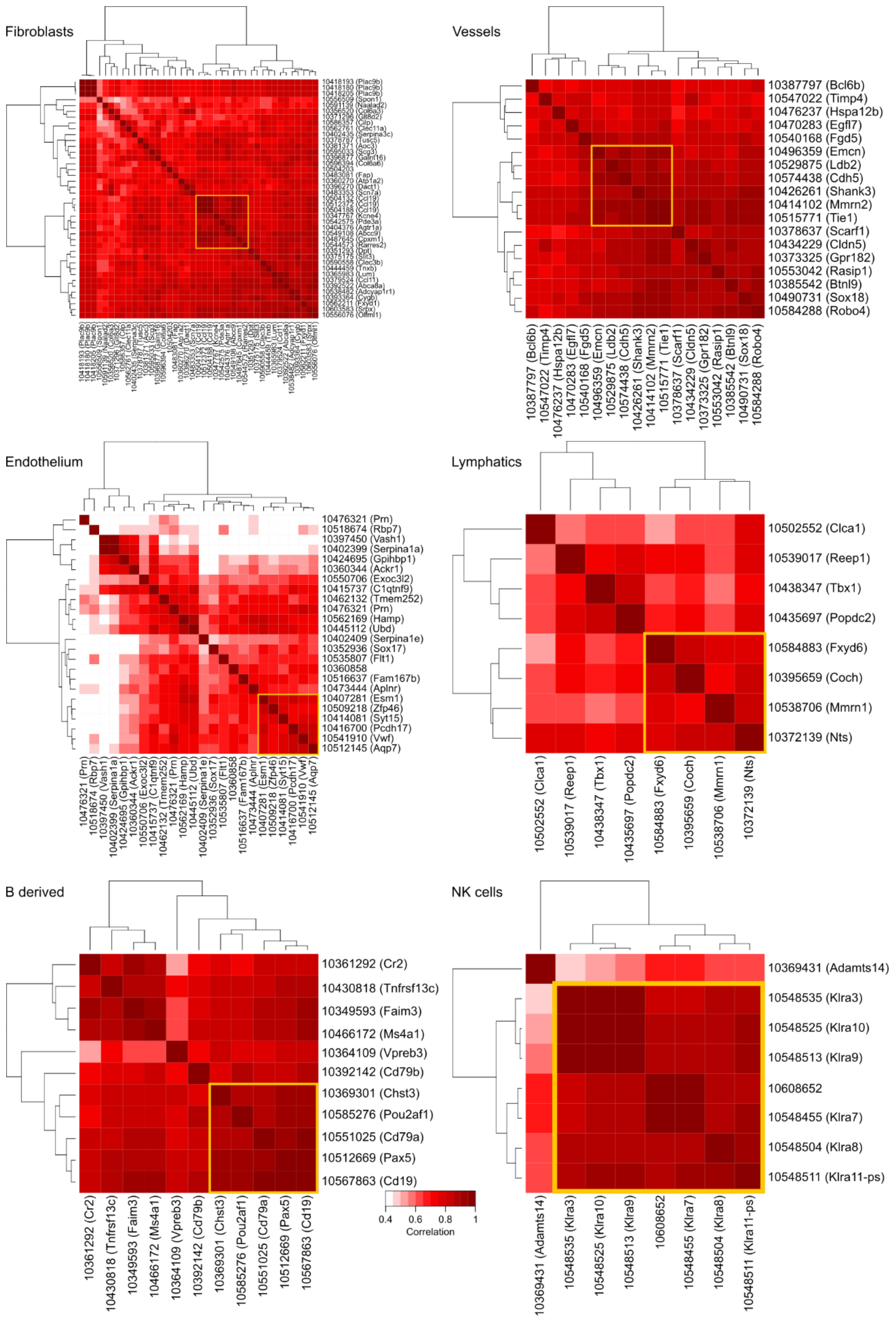


**Supplementary Figure 3: Selection of signatures for populations with numerous markers.**

Each heatmap is associated with a population that had 8 or more transcriptomic markers. The yellow rectangle indicates the chosen signature.

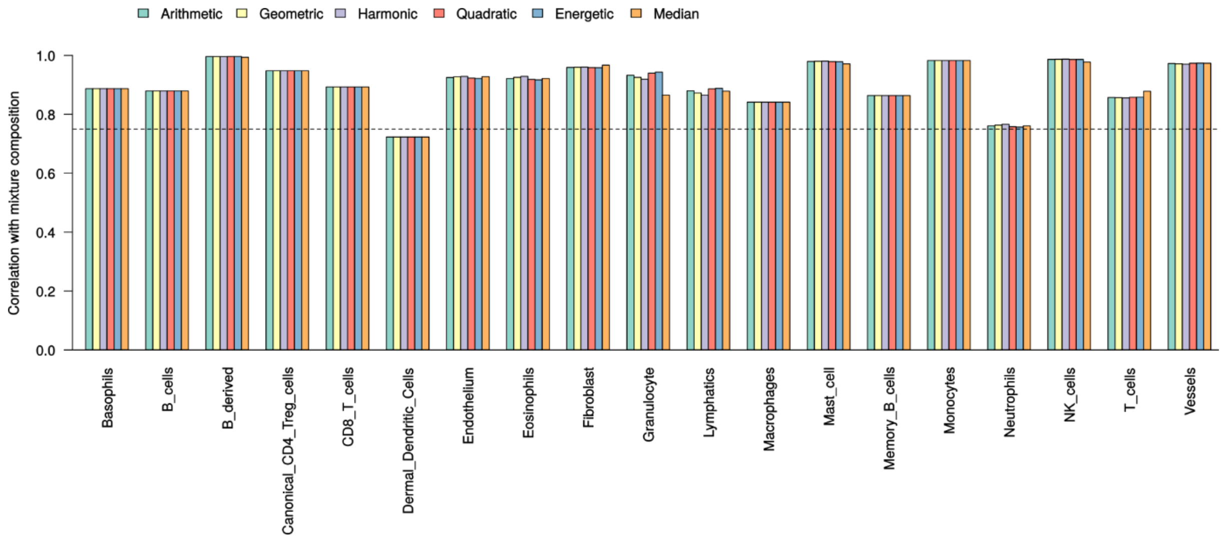


**Supplementary Figure 4: Comparison of different scoring methods.**

This figure relates to in silico simulated RNA mixtures. For each cell population, the correlation between the mixtures' compositions and the expression the of the signature was compared for various scoring methods.

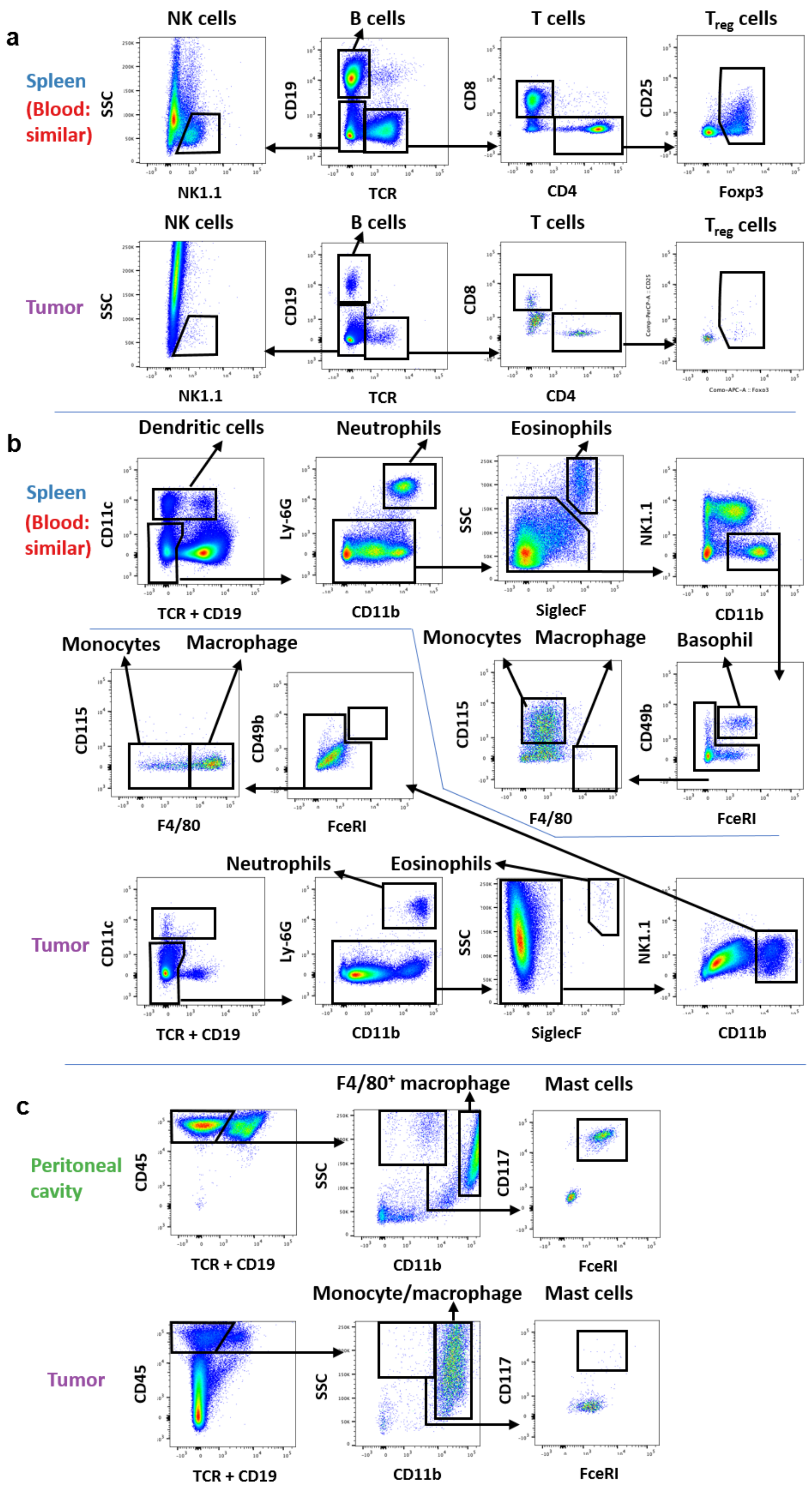
a
Spleen
(Blood: similar)
Tumor
NK cells
B cells
T cells
Treg cells
SSC
CD19
CD8
CD25
NK1.1
TCR
CD4
Foxp3
b
Dendritic cells
Neutrophils
Eosinophils
CD11c
Ly-6G
NK1.1
TCR + CD19
CD11b
SiglecF
Monocytes
Macrophage
Basophil
CD115
CD49b
F4/80
FceRI
c
Peritoneal cavity
F4/80+ macrophage
Mast cells
Monocyte/macrophage
CD45
CD117

22

**Supplementary Figure 5:** Representative gating strategy of flow cytometry analyses for **a.** T cells, B cells and NK cells; **b.** dendritic cells, granulocytes, monocytes and macrophages; and **c.** mast cells and peritoneal macrophages. Blood samples were not shown due to high similarity of the gating strategy to splenocytes

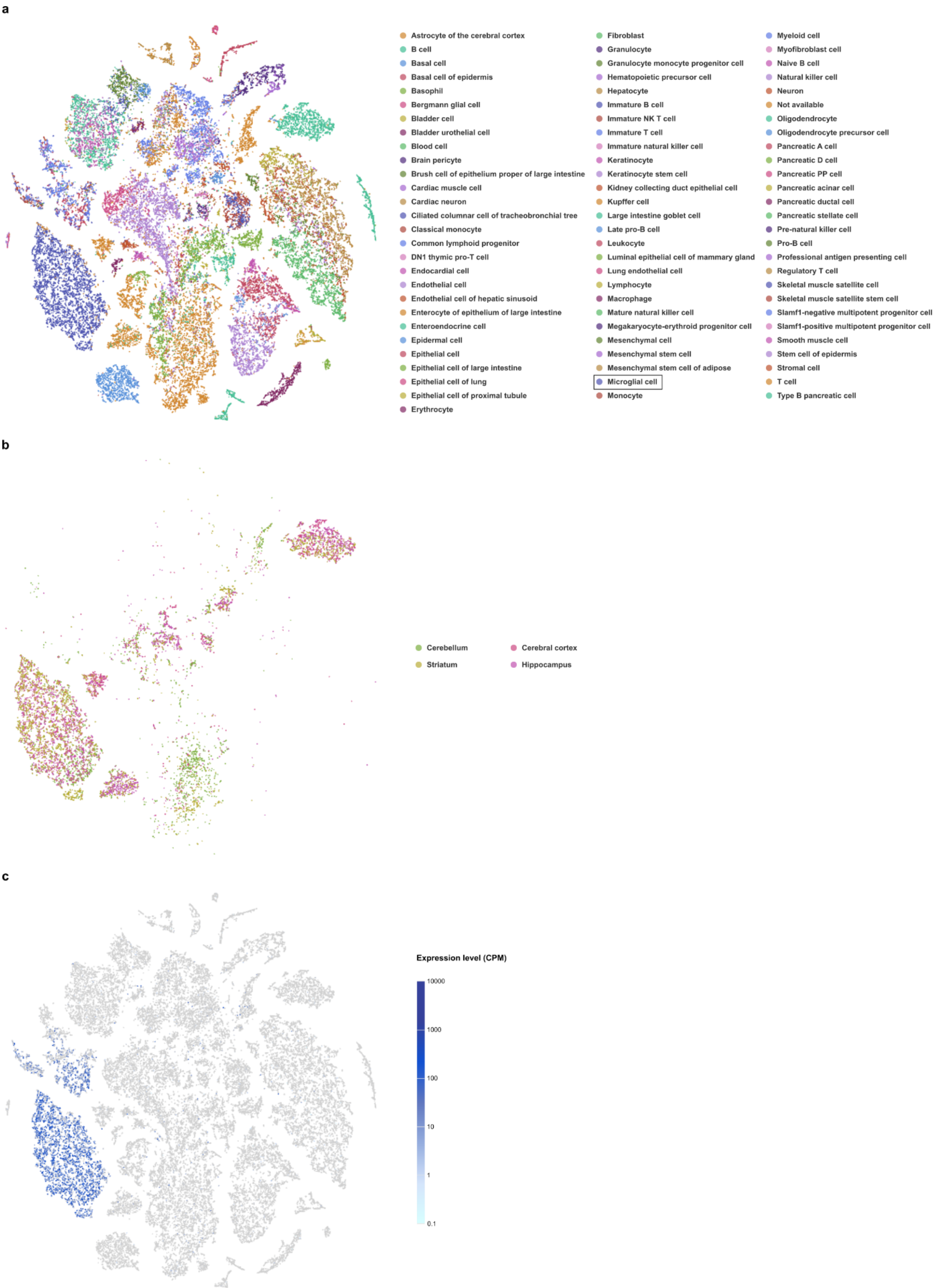


**Supplementary Figure 6: The mMCP-counter signature for monocytes / macrophages is also expressed in microglial cells in the brain**

This figure was generated using the Elixir platform from single-cell RNA-seq data from the Tabula Muris consortium.

**a.** t-SNE representation of the full dataset, colored by inferred cell type. This notably shows a cluster of microglial cells, dark blue, left of the plot.

**b.** Position of brain cells in the same t-SNE plot, colored by region of origin.

**c.** Expression level of the mMCP-counter signature for monocytes/macrophages, showing a strong expression in several cell clusters, including the microglial cell cluster.
